# The validated touch-video database

**DOI:** 10.1101/2023.10.31.565058

**Authors:** Sophie Smit, Anina N. Rich

## Abstract

Visually observing a touch quickly reveals who is being touched, how it might feel, and the broader social or emotional context, shaping our interpretation of such interactions. Investigating these dimensions is essential for understanding how tactile experiences are processed individually and how we empathise with observed sensations in others. Here, we expand available resources for studying visually perceived touch by providing a wide-ranging set of dynamic interactions that specifically focus on the sensory qualities of touch. The Validated Touch-Video Database (VTD) consists of a set of 90 videos depicting tactile interactions with a stationary left hand, viewed from a first-person perspective. In each video, a second hand makes contact either directly (e.g., with fingers or an open palm) or using an object (e.g., a soft brush or scissors), with variations across dimensions such as hedonic qualities, arousal, threat, touch type, and the object used. Validation by 350 participants (283 women, 66 men, 1 non-binary) involved categorising the videos as ‘neutral’, ‘pleasant’, ‘unpleasant’, or ‘painful’ and rating arousal and threat levels. Our findings reveal high inter-subject agreement, with painful touch videos eliciting the highest arousal and threat ratings, while neutral touch videos served as a baseline. Exploratory analyses revealed women rated the videos as more threatening and painful than men, suggesting potential gender differences in the visual perception of negatively valenced touch stimuli. The VTD provides a comprehensive resource for researchers investigating the sensory and emotional dimensions of observed touch.

## Introduction

Visually perceiving touch allows us to quickly gather important information about who is being touched (whether oneself or another person), the sensory qualities of the touch, and the broader emotional or social context (Smit, Ramírez-Haro, et al. 2024; Lee Masson and Isik 2023; Hertenstein et al. 2006). Much of the research on visual touch perception has focused on vicarious touch, where individuals report feeling tactile sensations in their own body simply by observing touch in others (for reviews see Gillmeister et al. 2017; Bufalari and Ionta 2013; Peled-Avron and Woolley 2022). This phenomenon is linked to the activation of the observer’s somatosensory cortex (Bufalari et al. 2007; Rigato et al. 2019; Bolognini et al. 2011; 2013; Blakemore et al. 2005), suggesting that we may simulate observed touch as though we are experiencing it ourselves (Decety and Jackson 2004; Gallese, Keysers, and Rizzolatti 2004; Keysers and Gazzola 2009; Smit et al. 2023). Given the importance of visual touch perception in interpreting sensory and emotional experiences in ourselves and others, we need to fully understand how we process detailed sensory qualities of observed touch, as well as emotional-affective dimensions like arousal and threat.

Different aspects have been shown to influence how observed touch is perceived and processed. For instance, stimuli depicting more threatening or painful experiences elicit stronger vicarious responses compared to neutral or non-threatening touch (Holle et al. 2011; Ward, Schnakenberg, and Banissy 2018; Li and Jamie Ward 2022; Smit, Crossley, et al. 2024). Physical factors, such as the body part touched, congruent posture, viewing perspective, and the realism of the stimuli, also play an important role in these processes (Holle et al. 2011; Medina and DePasquale 2017). However, because many studies use stimuli that differ across multiple dimensions simultaneously, it is difficult to isolate the specific influence of these factors. Research using controlled stimuli would enhance the potential for systematic investigations into how elements like arousal, threat, and hedonic quality shape responses to observed touch.

Research on visual touch perception has greatly enriched our understanding of somatosensory processes (Rigato et al. 2019; Adler et al. 2016; Ward, Schnakenberg, and Banissy 2018; Bufalari et al. 2007; Walker et al. 2017; Smit et al. 2023; Bolognini et al. 2011; Smit, Rich, and Zopf 2019). To date, however, relatively little attention has been given to the dynamic and detailed sensory qualities of touch. Although excellent databases such as the Social Touch Picture Set (SToPS) (Schirmer et al. 2015) and the Socio-Affective Touch Expression Database (SATED) (Masson and Beeck 2018) have advanced our understanding of the social aspects of observed touch, few resources offer such systematically controlled and validated sensory-focused stimuli. To address this gap, we introduce the Validated Touch-Video Database (VTD), a standardised set of 90 videos capturing a wide range of close-up tactile interactions. These videos vary in key dimensions—hedonic quality, arousal, and threat—while ensuring consistency in factors such as the body parts involved (e.g., a right hand interacting with a left hand from a first-person perspective) and the visual context. The VTD provides a robust resource for advancing our exploration of the neural and psychological mechanisms underlying complex and dynamic visual touch processing.

To validate the VTD, 350 participants categorised each video as neutral, pleasant, unpleasant, or painful and rated the strength of the hedonic quality. Participants also assessed perceived levels of arousal and threat for each video. These dimensions were chosen based on their fundamental role in shaping touch perception. Hedonic quality is a key determinant of affective responses to touch and has been shown to influence both neural and behavioural reactions (McGlone, Wessberg, and Olausson 2014; Olausson et al. 2016). It reflects the degree of pleasantness or unpleasantness associated with a touch experience, encompassing both rewarding and aversive aspects. Pain, as an intensely aversive sensation, represents the negative extreme of hedonic quality, while gentle caresses or soft strokes represent the positive extreme (McGlone, Wessberg, and Olausson 2014). Subjective arousal reflects the perceived intensity of the sensory and emotional response to touch, and threat perception plays a crucial role in distinguishing between safe and potentially harmful touch, both affecting attentional engagement and physiological reactivity (Vogt et al. 2008; Abra, Mirams, and Fairhurst 2024; Koster et al. 2004; Poliakoff et al. 2007). By systematically varying and validating these dimensions, the VTD provides a unique resource for fine-grained investigation of how people visually perceive touch, enabling researchers to explore the interplay between sensory details and emotional-affective processing.

As gender differences in vicarious sensory perception have been observed, particularly in neural responses to observed pain (Grice-Jackson et al. 2017; Singer et al. 2006; Li and Jamie Ward 2022; Yang et al. 2009; Smit, Crossley, et al. 2024), we included an exploratory analysis to test whether men and women evaluated the observed touch itself differently. While a previous study found no gender differences in how valence, arousal, and naturalness were rated for observed touch in broader social contexts (Masson and Beeck 2018), such differences may emerge when assessing the perception of more detailed sensory qualities, particularly in painful touch contexts. Examining how men and women rate various dimensions of detailed visual touch stimuli could reveal differences at the initial stage of stimulus evaluation.

Our analysis demonstrated very strong consistency in subjective arousal and threat ratings between participants, with substantial agreement in categorising the hedonic qualities of the videos. When comparing genders, men were more likely to categorise the observed touch as ‘neutral’, while women more often classified the touch as ‘painful’ and rated painful videos as more intense. Additionally, women rated the videos as more threatening overall, though no gender differences were observed in arousal ratings. These findings suggest that women may exhibit heightened sensitivity to touch stimuli with negative valence, which aligns with prior research on vicarious responses to pain. All videos, along with validation data, are freely accessible online, providing an open resource for future research on the sensory and emotional dimensions of touch.

## Materials and methods

### Data, code & video availability

**OSF**: Videos, analysis code, and validation data https://osf.io/jvkqa/

**YouTube**: Video playlist https://bit.ly/YouTubeVTD

**GitHub**: Supplementary materials https://sophiesmit1.github.io/VTD/

### Creation of the videos

The aim of this study was to create and validate a set of close-up touch videos that capture a wide range of tactile interactions to a hand, varying systematically across the dimensions of hedonic quality, arousal, and threat. We produced 90 short videos, each featuring a Caucasian female hand placed on a dark background, with the hand always viewed from a first-person perspective, palm facing down (see Fig. 1). The videos varied in duration from 1.4 seconds to 11.8 seconds (*M* = 3.8 seconds).

**Figure 1.**
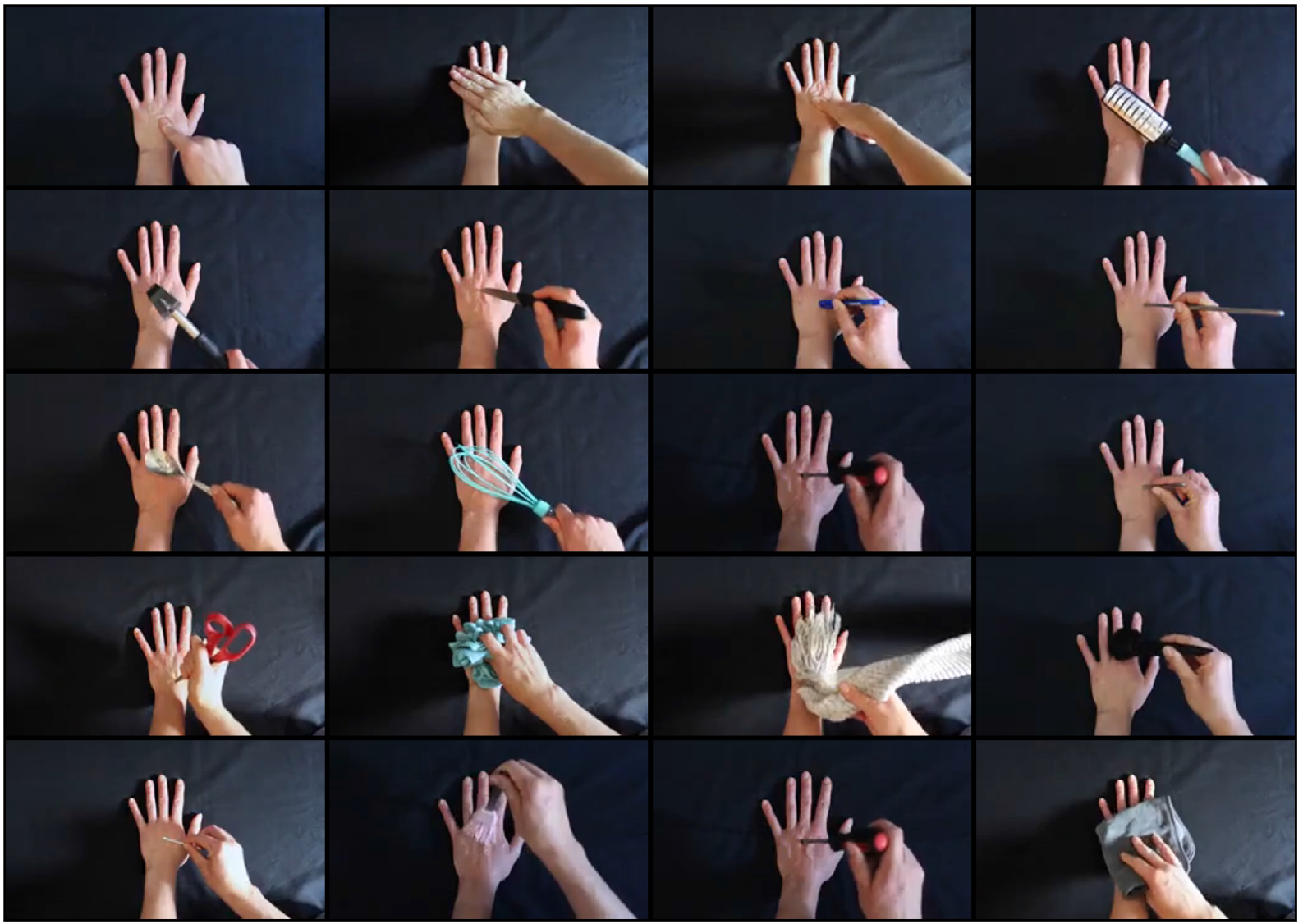
Still frames for a subset of videos from the Validated Touch-Video Database. These frames illustrate a variety of touch interactions captured from a first-person perspective, depicting different hedonic qualities including neutral, pleasant, unpleasant, and painful touch. The database includes 90 systematically controlled videos that vary across dimensions such as arousal, threat, and touch type, providing a comprehensive resource for investigating the emotional and sensory dimensions of observed touch.

In each video, a second hand touches the stationary hand, either directly (e.g., using fingers or a flat hand) or using an object (e.g., a soft brush, scissors). To create neutral videos, we depicted gentle touches with minimal pressure using non-threatening objects or directly skin contact with another hand. The pleasant videos show slow, stroking movements across the hand using either a soft brush or fingers, aiming to evoke a positive, comforting sensation. Unpleasant and painful videos show aversive interactions, such as touching the hand with a syringe or pinching the skin with tweezers. These interactions appeared realistic enough to suggest discomfort or pain without causing any actual harm or distress to the model. We carefully constructed each video to represent a distinct tactile experience while maintaining consistency in presentation. The dark background and consistent camera angle helped focus attention on the tactile interactions, minimising external distractions.

### Validation of the videos

#### Participants

Participants were recruited via the Macquarie University subject pools and consisted of undergraduate students naïve to the purpose of the study (*N* = 350; 283 women, 66 men; 1 non-binary, *M* = 24 years, *SD* = 8.66 years, range = 17-63 years). As only one non-binary participant was included, we excluded their data from gender-based analyses.

We asked participants to evaluate the videos based on hedonic qualities, perceived arousal, and threat. Due to the large number of videos, some of which were unpleasant to watch, we set up two separate questionnaires with 45 videos each, each questionnaire comprising of an equal number of neutral, pleasant, unpleasant, and painful videos. Each questionnaire took approximately 15 minutes to complete. Participants were free to decide whether they wanted to participate in one or both questionnaires and received course credit accordingly. The study was approved by the Macquarie University Human Research Ethics committee and participants provided written consent.

#### Experimental procedure

Participants completed the questionnaire remotely via the Qualtrics online platform. Participants were instructed to complete the study on a desktop or laptop to ensure proper video presentation and avoid display size variability. The videos were presented in a random order, and auto-play was disabled to ensure participant readiness and engagement. Each video began with a white screen displaying the instruction, “Press play to start video”, requiring participants to actively initiate the onset. After watching each video, participants categorised the touch based on its hedonic quality, then rated the strength of the chosen category, perceived threat, and arousal, all on a 1-10 scale (from not at all to extremely). The following four questions were presented after each video:

- **Q1**: How would you categorise the touch in this video? [Options: neutral, pleasant, unpleasant, painful]
- **Q2**: How [pleasant/unpleasant/painful] (based on the previous response) was the touch?
- **Q3**: How threatening was the touch?
- **Q4**: How arousing was this video? (Arousal in terms of a feeling, emotion, or response)

#### Analyses

In this study, we used Bayesian statistical methods for all analyses, including t-tests, ANOVAs, correlation assessments, and contingency tables. The analyses were conducted using the R programming environment (R Core Team 2023), with the Bayes Factor R package (Morey, Rouder, and Jamil 2018) to calculate Bayes factors (BF). We adopted a default prior based on a Cauchy distribution centred at zero with a scale parameter of *r* = 0.707, allowing for a range of possible effect sizes. Bayes factors quantify the relative strength of evidence for the alternative hypothesis compared to the null hypothesis. For instance, a Bayes factor of 3 indicates that the evidence from our data is three times stronger for the alternative hypothesis than for the null hypothesis. Generally, a Bayes factor greater than 1 indicates support for the alternative hypothesis, while a factor less than 1 favours the null hypothesis, and values between 1 and 3 are considered insufficient for making a definitive conclusion (Jeffreys and Jeffreys 1998; Dienes 2011; Morey, Romeijn, and Rouder 2016; Rouder et al. 2009).

We assessed the consistency of participant ratings for arousal and threat across the 90 videos in our database using the Intraclass Correlation Coefficient (ICC). Specifically, we used a two-way mixed-effects model (ICC3k) to calculate the mean-rating consistency, providing a robust measure of agreement (Koo and Li 2016). ICC estimates were computed along with 95% confidence intervals using the R package Psych (Revelle 2024). A high ICC value indicates strong inter-rater reliability, confirming consistency in participant ratings across videos.

We used Fleiss’ Kappa (Landis and Koch 1977) to assess the consistency of participants’ categorisation of the videos into four distinct hedonic categories: neutral, pleasant, unpleasant, and painful. For each of the 90 videos, we calculated the proportion of participants who classified the video into each of the four categories. Fleiss’ Kappa values range from -1 to 1 and provide a scale for interpreting the strength of agreement among participants. Negative values suggest less agreement than expected by chance, 0 indicates chance-level agreement, and positive values indicate better agreement than chance. Fleiss’ Kappa values can be interpreted as follows: < 0 poor; 0 - 0.20 slight; 0.21 - 0.40 fair; 0.41 - 0.60 moderate; 0.61 - 0.80 substantial; 0.81 - 1 almost perfect agreement (Landis and Koch 1977).

Recognising potential overlap in hedonic categorisation—where some videos may be classified as unpleasant by certain participants and painful by others—we also conducted a focused analysis on a subset comprising the top 10 videos from each category (40 videos total). This subset was selected to provide clear distinctions between hedonic categories, ensuring that the hedonic quality of touch is well-defined for subsequent research. We also examined the arousal and threat ratings for this clearly categorised subset to gain a deeper understanding of how these dimensions vary across different hedonic qualities.

To explore potential gender-specific differences in the assessment of our video stimuli, we examined the dimensions of arousal, threat, and hedonic qualities across all 90 videos, stratified by participant gender. While unbalanced samples may present challenges for frequentist approaches, Bayesian methods address these issues by directly estimating the likelihood of the observed effects and incorporating the uncertainty caused by unequal group sizes (Morey, Romeijn, and Rouder 2016; Rouder et al. 2009; Dienes 2011; Kruschke 2013).

## Results

### High agreement across participants for arousal and threat ratings

The ICC analysis yielded high values for both arousal and threat ratings, suggesting a strong level of agreement among participants’ ratings (values greater than 0.81 indicate almost perfect agreement; Koo and Li, 2016). Across all participants, the ICC values were very high for both arousal (*ICC* = 0.98, 95% *CI*: 0.98 - 0.99) and threat (*ICC* = 0.99, 95% *CI*: 0.99 - 1.00). These findings demonstrate strong consistency in participants’ arousal and threat ratings across the video stimuli. Overall, women provided higher threat ratings (*M* = 3.50) than men (*M* = 2.99; *BF* > 1000), whereas arousal ratings did not differ between genders (women: *M* = 3.83, men: *M* = 3.76; *BF* = 0.05; see Fig. 2).

**Figure 2.**
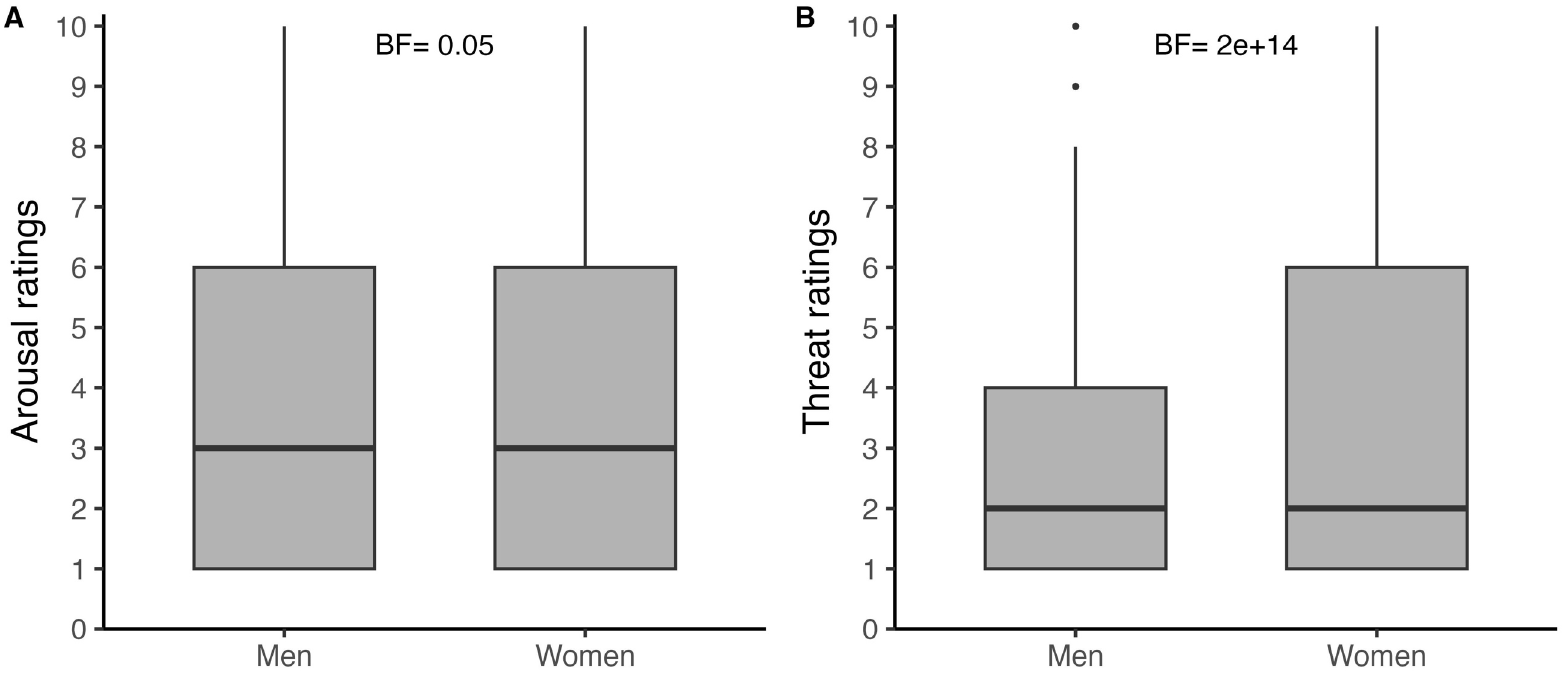
Arousal and threat ratings by gender. The box plots display the median ratings for arousal and threat across all 90 videos, separated by gender. Participants rated the touch on a scale from 1 (not at all) to 10 (extremely), with a total of 283 women and 66 men.

### Fair agreement across participants for hedonic categorisation

Most videos were categorised by participants as neutral (35%), followed by unpleasant (33%), pleasant (17%) and painful (14%). We assessed the inter-rater reliability for categorising the 90 videos into four hedonic categories: neutral, pleasant, unpleasant, or painful. The Fleiss’ Kappa value for the full set of videos was 0.274, indicating ‘fair’ agreement. The lower reliability reflects overlap in categorisation, with some videos being classified as equally neutral/pleasant or unpleasant/painful (see supplementary materials: https://sophiesmit1.github.io/VTD/). A Bayesian contingency table showed an overall difference in how the genders categorised the videos (*BF* > 1000) with a higher percentage of men categorising videos as neutral (*BF* = 884.9) and a higher percentage of women categorising videos as painful (*BF* > 1000) (see Fig. 3).

**Figure 3.**
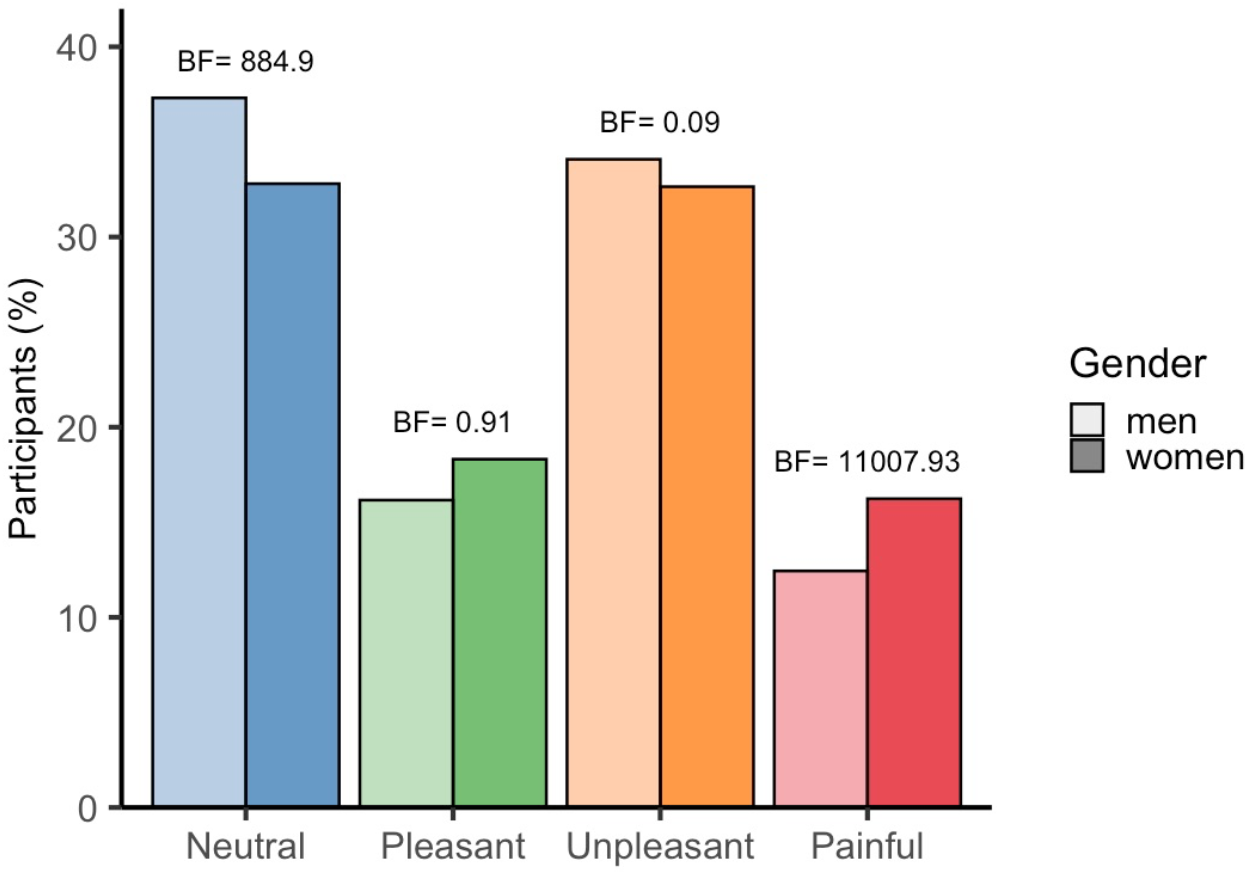
Hedonic categorisation of touch videos by gender. The bar graphs display the categorisation of all 90 videos into one of four hedonics categories, separated by gender.

To provide researchers with a subset of videos that clearly depict one of the hedonic categories, we selected the top 10 videos from each category—those with the highest percentage of participants agreeing on their categorisation—creating a 40-video subset. Agreement levels were as follows: neutral videos (70% to 85%), pleasant videos (63% to 91%), unpleasant videos (58% to 70%), and painful videos (45% to 83%) (see Table 1). For example, in the neutral category, the most clearly categorised video had 85% agreement, while the 10th video had 70% agreement. This indicates consistent categorisation and clear distinctions in hedonic qualities within this subset of 40 videos, with only some of the painful videos also being categorised as unpleasant. For this subset, the Fleiss’ Kappa value increased to 0.387.

**Table 1.** Percentage of sample agreement in categorising top 10 videos by hedonic quality, with corresponding mean arousal and threat ratings for these videos; 1-10 scale (not at all - extremely).

| Top 10 | Description (neutral videos) | Neutral (%) | Pleasant (%) | Unpleasant (%) | Painful (%) | Threat (1-10) | Arousal (1-10) |
| --- | --- | --- | --- | --- | --- | --- | --- |
| 1 | single touch with finger | 85 | 6 | 8 | 1 | 1.4 | 1.8 |
| 2 | single touch with pen | 79 | 4 | 17 | 0 | 4.2 | 3.6 |
| 3 | single push with a spoon | 79 | 7 | 14 | 0 | 3.2 | 3.7 |
| 4 | single touch with cotton bud | 78 | 19 | 2 | 1 | 1.3 | 4.5 |
| 5 | repeated tap with finger | 77 | 11 | 12 | 0 | 2.6 | 3.0 |
| 6 | single touch with flat hand | 75 | 14 | 11 | 0 | 5.2 | 4.0 |
| 7 | single touch with plastic brush | 75 | 23 | 2 | 0 | 1.3 | 4.2 |
| 8 | repeated touch with cotton bud | 73 | 20 | 6 | 1 | 1.4 | 2.7 |
| 9 | single push with a plastic whisk | 71 | 10 | 19 | 1 | 6.5 | 5.3 |
| 10 | Repeated touch with finger | 70 | 6 | 24 | 0 | 1.8 | 2.2 |
| Top 10 | Description (pleasant videos) | Neutral | Pleasant | Unpleasant | Painful | Threat | Arousal |
| 1 | long stroke with soft brush all over the hand | 6 | 91 | 3 | 0 | 1.5 | 3.0 |
| 2 | single stroke top to bottom with soft brush | 8 | 88 | 4 | 0 | 3.5 | 3.7 |
| 3 | stroke top to bottom with shawl | 12 | 81 | 4 | 2 | 2.6 | 2.8 |
| 4 | single stroke top to bottom with plastic brush | 17 | 81 | 2 | 0 | 1.4 | 3.4 |
| 5 | stroke with soft sock | 11 | 79 | 10 | 0 | 3.5 | 3.8 |
| 6 | repeated stroke with plastic brush | 15 | 79 | 5 | 1 | 1.2 | 2.2 |
| 7 | single stroke top to bottom with cotton pad | 26 | 73 | 1 | 1 | 1.4 | 3.8 |
| 8 | (long) stroke with multiple fingers all over the hand | 18 | 71 | 10 | 1 | 6.6 | 5.6 |
| 9 | stroke top to bottom with sponge | 33 | 63 | 3 | 1 | 2.6 | 4.0 |
| 10 | touch top to bottom with fabric | 33 | 63 | 3 | 1 | 1.3 | 2.1 |
| **Top 10** | **Description (unpleasant videos)** | **Neutral** | **Pleasant** | **Unpleasant** | **Painful** | **Threat** | **Arousal** |
| 1 | touch top to bottom with scissors | 6 | 6 | 70 | 18 | 4.8 | 4.2 |
| 2 | repeated touch with pencil | 22 | 1 | 67 | 10 | 4.7 | 4.7 |
| 3 | touch top to bottom with hammer | 30 | 4 | 63 | 3 | 3.3 | 3.1 |
| 4 | hand grabbing wrist | 12 | 6 | 61 | 21 | 3.5 | 3.3 |
| 5 | repeated touch with metal chopstick | 23 | 5 | 59 | 12 | 2.4 | 2.6 |
| 6 | scratching with fingernails | 18 | 17 | 59 | 6 | 1.8 | 3.2 |
| 7 | stroke top to bottom with stanley knife | 6 | 3 | 59 | 32 | 4.5 | 5.0 |
| 8 | repeated touch with rolling pin | 20 | 2 | 58 | 20 | 8.0 | 6.7 |
| 9 | single touch with screwdriver | 11 | 0 | 58 | 31 | 2.5 | 2.7 |
| 10 | touch top to bottom with nail file | 8 | 6 | 58 | 28 | 1.4 | 2.6 |
| **Top 10** | **Description (painful videos)** | **Neutral** | **Pleasant** | **Unpleasant** | **Painful** | **Threat** | **Arousal** |
| 1 | single stab with nail file | 2 | 1 | 14 | 83 | 1.2 | 4.7 |
| 2 | single stab with scissors | 1 | 2 | 14 | 83 | 2.9 | 3.6 |
| 3 | scratch top to bottom with scissors | 3 | 1 | 31 | 65 | 1.5 | 4.1 |
| 4 | single pinch with tweezers | 2 | 2 | 33 | 63 | 6.0 | 4.9 |
| 5 | repeated touch with nail file | 2 | 1 | 36 | 61 | 1.4 | 2.4 |
| 6 | single injection with sharp needle | 3 | 0 | 38 | 59 | 5.2 | 4.7 |
| 7 | repeated touch with nail file | 5 | 1 | 46 | 49 | 5.9 | 4.6 |
| 8 | repeated punch with hand | 8 | 3 | 42 | 46 | 1.5 | 3.3 |
| 9 | single punch with hand | 10 | 1 | 44 | 45 | 1.5 | 4.0 |
| 10 | repeated touch with hammer | 9 | 0 | 46 | 45 | 5.1 | 4.3 |

### Strength of hedonic quality

For the subset of 40 videos based on the highest hedonic categorisation, we assessed how intense participants perceived the hedonics of the touch to be (see Fig. 4). We focused on non-neutral touch videos (pleasant, unpleasant, or painful) and asked participants, “How pleasant/unpleasant/painful was the touch?” A Bayesian ANOVA showed a difference in ratings across categories (*BF* > 1000), with the highest ratings for pleasant videos (*M* = 4.38), followed by painful videos (4.20) and unpleasant videos (3.10). Bayes factors indicated strong evidence for a difference in intensity between unpleasant and both pleasant (*BF* > 1000) and painful videos (*BF* > 1000). However, there was moderate evidence suggesting no difference in intensity between pleasant and painful videos (*BF* = 0.12), indicating they were perceived as similarly intense. Women rated the painful videos as more painful than men (*BF* = 12.36) (see Fig. 5).

**Figure 4.**
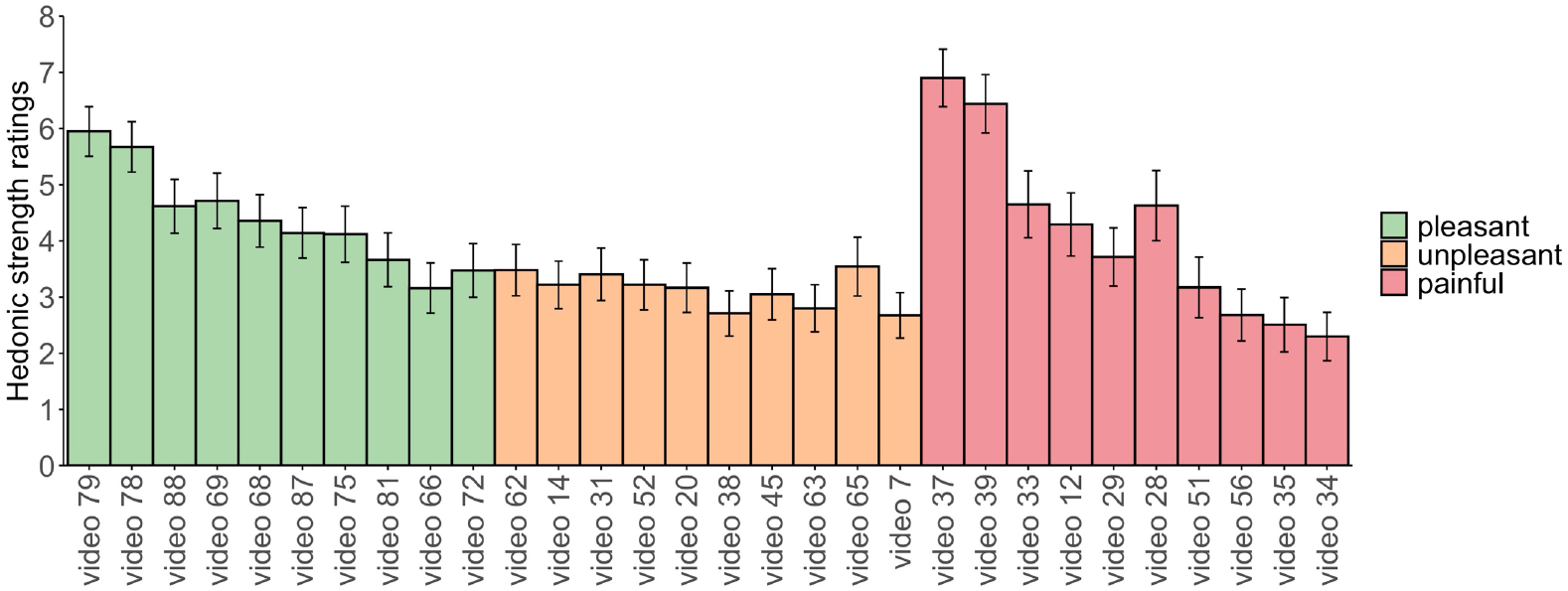
Hedonic quality strength ratings for the top 10 videos in each category. Each bar graph displays the average hedonic strength ratings for the entire sample, accompanied by 95% confidence intervals. Participants rated the touch on a scale from 1 (not at all) to 10 (extremely). The top 10 most distinctly categorised videos are organised from left to right, with pleasant (green), unpleasant (orange), and painful (red).

**Figure 5.**
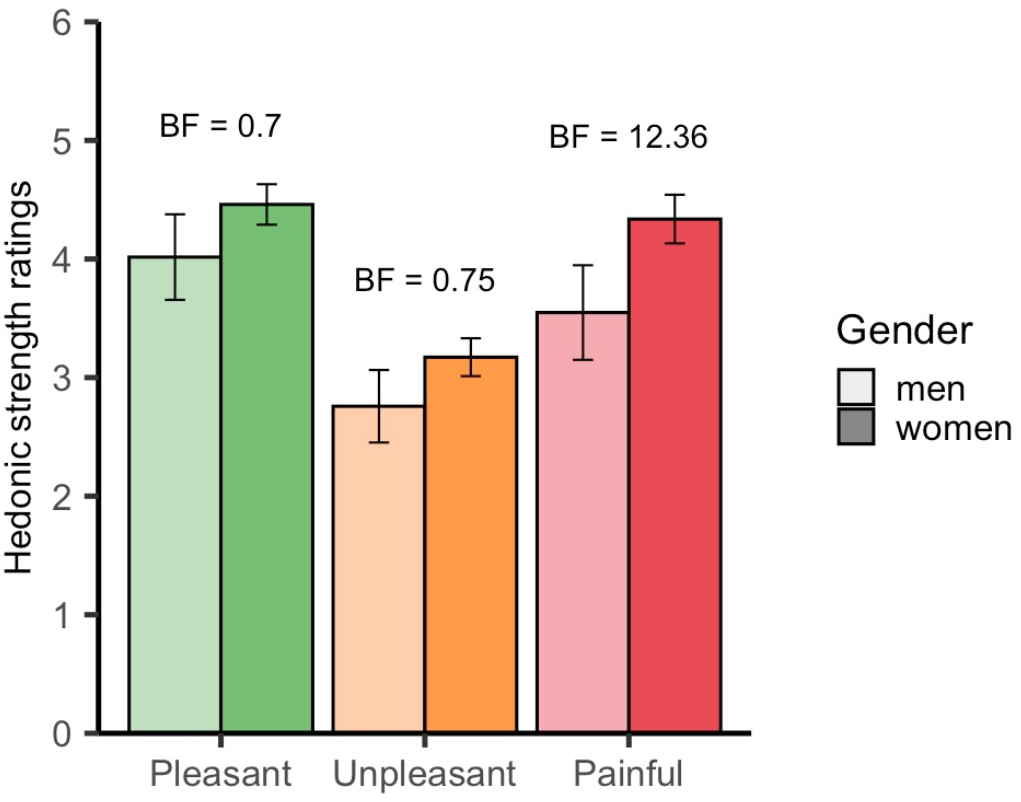
Hedonic strength ratings for each category divided by gender. Each bar shows the average hedonic strength ratings for a subset of videos (top 10 per category), differentiated by gender (light bars for men and dark bars for women) with 95% confidence intervals. Videos are grouped as pleasant (green), unpleasant (orange), and painful (red).

### Correlation between hedonic qualities and arousal and threat ratings

Our goal was to create a database of videos that depicted touch with varying hedonic qualities and substantial variety in arousal levels. Ideally this would allow for a selection of videos that differ in whether they are pleasant or unpleasant while being matched on arousal. We also aimed to include videos with a range of threat levels, as this remains an underexplored dimension of visual touch perception. Although arousal and threat are often correlated, pleasant videos would likely be rated as highly arousing yet low in threat. Additionally, we aimed to include videos perceived as truly neutral—neither pleasant, unpleasant, nor painful— while also being low in arousal and threat. To evaluate whether we achieved this, we examined the interplay between the hedonic qualities of touch and participants’ ratings for arousal and threat (see Fig. 6, including Bayes factors for pairwise comparisons). Bayesian ANOVAs confirmed differences in arousal (*BF* > 1000) and threat (*BF* > 1000) ratings across the hedonic categories. Pleasant and unpleasant videos were rated similarly in arousal (*M* = 4.06 vs. *M* = 4.09), whereas painful videos received the highest arousal ratings (*M* = 5.50). Painful videos were most threatening (*M* = 6.35) followed by unpleasant videos (*M* = 4.29), both with substantial variability within the hedonic categories (unpleasant range = 2.45 - 6.58; painful range = 4.54 - 7.93). Neutral videos had the lowest arousal ratings (*M* = 2.06), and both neutral and pleasant videos were rated similarly low in threat (*M* = 1.53 vs. *M* = 1.39).

**Figure 6.**
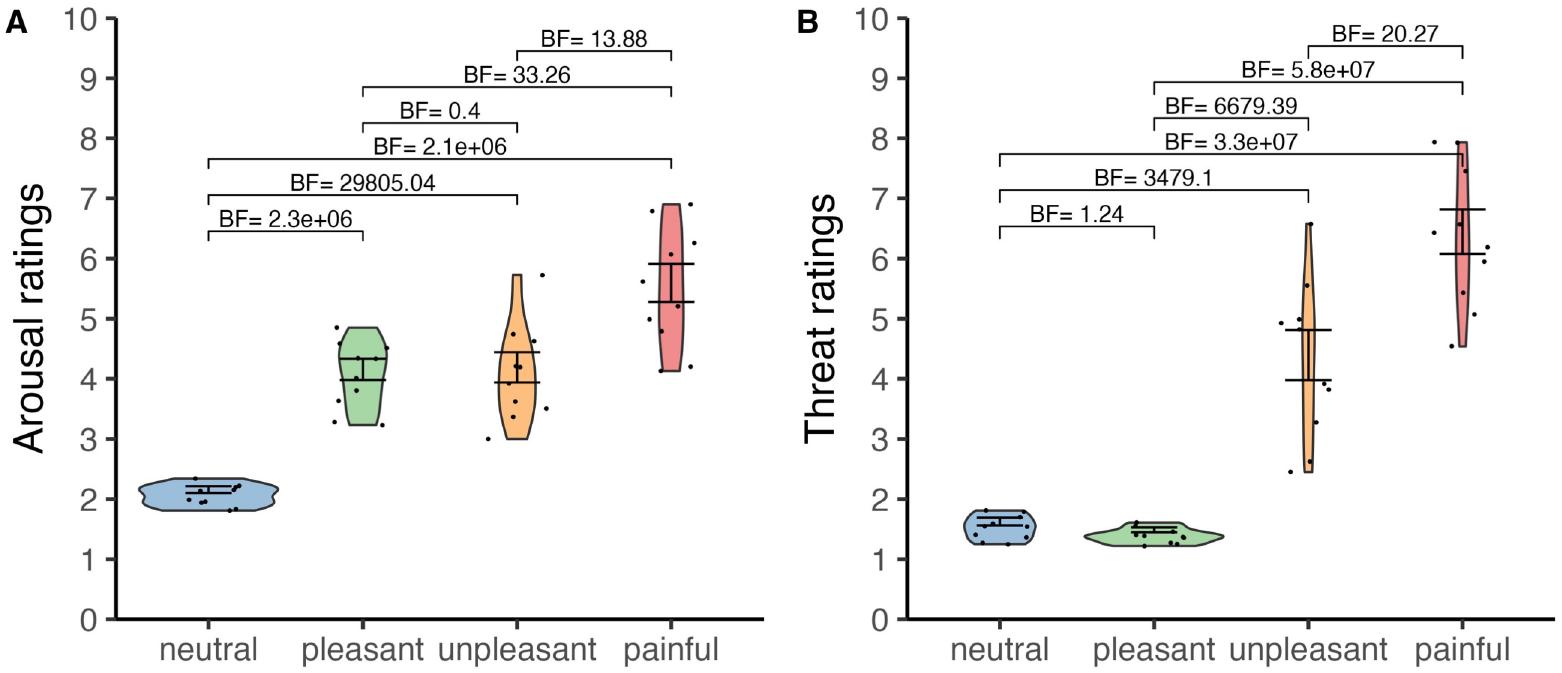
Average ratings for arousal and threat across the top 10 videos in each hedonic category. Participants rated each video on a scale from 1 (not at all) to 10 (extremely). Scatter plots within the violins represent mean scores for individual videos.

### Videos & validation data

To facilitate research use, we compiled a summary table that includes a description of each video alongside its corresponding YouTube link (https://sophiesmit1.github.io/VTD/). Within this table, users can sort the collection of 90 videos based on key metrics—such as hedonic quality, threat, and arousal—either ascending or descending (see Fig. 7). It also allows users to find videos that align with specific keywords or criteria. The table is also downloadable in multiple formats, including CSV, Excel, and PDF. This feature allows users to efficiently filter and select videos based on specific characteristics of interest.

**Figure 7.**
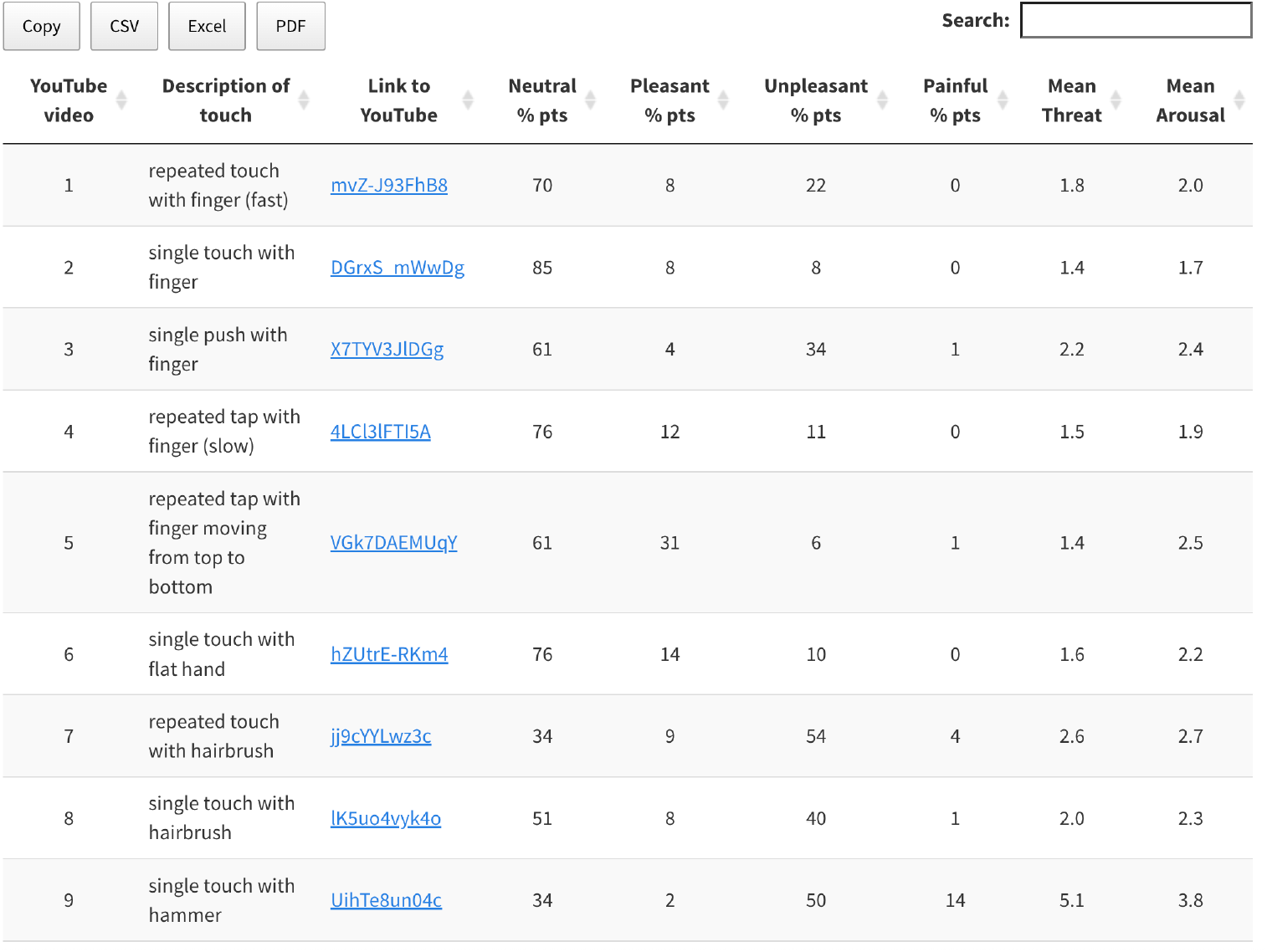
Overview of the videos along with their validation data. Preview of the overview table which can be found on GitHub (https://sophiesmit1.github.io/VTD/). The 90 videos in this table can be arranged based on the highest or lowest values for each index, such as hedonic quality (percentage of participants), threat, and arousal (1 = not at all, 10 = extremely).

## Discussion

The aim of this study was to create a systematically defined database of videos depicting a diverse range of tactile interactions to a hand, enabling researchers to investigate the nuances of visual touch perception while controlling for specific dimensions. High inter-subject agreement confirmed that the videos effectively captured varying levels of subjective arousal and threat. A subset of 40 videos was reliably categorised into neutral, pleasant, unpleasant, and painful touch, with differences observed in perceived arousal and threat levels across these categories. Videos depicting very painful or pleasant touch were associated with the highest arousal ratings, reflecting the previously documented relationship between valence and arousal (Bradley and Lang 1994; Masson and Beeck 2018). The inclusion of a neutral category in the database allows researchers to draw meaningful comparisons between emotionally charged touch interactions and neutral touch.

Our findings reveal gender-related differences in how tactile interactions are perceived. Although both genders provided similar arousal ratings, women rated the videos as more threatening and were more likely to classify them as painful, while men tended to categorise them as neutral. Women also reported higher intensity ratings for painful videos. These findings suggest that women may exhibit greater sensitivity to negatively valenced visual touch stimuli, consistent with prior research on vicarious responses to pain (Grice-Jackson et al. 2017; Singer et al. 2006; Li and Jamie Ward 2022; Yang et al. 2009; Smit, Crossley, et al. 2024). Notably, an earlier study reported no gender differences in ratings for valence, arousal, and naturalness of observed touch in broader social contexts (Masson and Beeck 2018), though the sample size was small. In contrast, our study, with a large sample size, reveals that gender differences emerge when participants assess detailed touch videos, particularly in negative contexts. These results suggest it may be useful to consider gender as a factor when evaluating responses to visually perceived touch and pain stimuli.

The VTD includes a broad range of tactile interactions, providing a valuable tool for examining how different types of touch affect individuals prone to vicarious tactile sensations. Most research on vicarious sensory perception has focused on reactions to observed neutral or threatening touch and pain (Giummarra et al., 2015; Grice-Jackson et al., 2017; Osborn and Derbyshire, 2010; Vandenbroucke et al., 2013; Ward et al., 2018; Ward and Li, 2022), with relatively little attention to pleasant touch. While fMRI studies have deepened our understanding of the neural correlates of empathy for pleasant and unpleasant sensations (Ebisch et al., 2011; Lamm et al., 2015; Morrison et al., 2011; Riva et al., 2018), there remains a need to explore subjective experiences associated with observing touch across different hedonic qualities (e.g., Smit, Crossley, et al. 2024). A recent review identified soft materials and slow, gentle stroking as key factors eliciting pleasurable sensations during direct touch (Taneja et al., 2021). Here, we incorporated such elements to create pleasant visual touch stimuli, which can be used in research settings ranging from therapeutic applications of vicarious touch (Giummarra et al., 2016; Makary et al., 2018; Ramachandran and Altschuler, 2009; Ramachandran and Rodgers-Ramachandran, 1996) to digital marketing and virtual environments (Luangrath et al., 2022). Access to a collection of video stimuli showing diverse touch interactions, including both pleasant and unpleasant touch, allows for a fuller understanding of vicarious tactile experiences.

Videos in the database differ in duration, with pleasant videos often featuring slower, longer strokes, while neutral videos displayed brief, simple touches. Researchers may wish to modify these videos in terms of duration, presentation size, or orientation to suit their study’s needs. For instance, in studies examining time-locked responses to observed touch using electroencephalography (EEG), it may be beneficial to standardise the video length. Some studies may also require the hand to be presented from a first- or third-person perspective to suggest touch to oneself or another. In a separate study (Smit, Ramírez-Haro, et al. 2024), we shortened and standardised the videos to 600 milliseconds, presented them in a smaller size, and varied the orientation (flipped horizontally, vertically, or both). Results from an independent participant sample revealed strong correlations between ratings for the original and modified videos across valence, arousal, and threat (mean across modifications: *r* = 0.89), indicating that ratings remain robust even when stimuli are adapted. Both the original and adapted videos are available online (https://osf.io/jvkqa/).

Our participant pool consisted primarily of psychology undergraduate students, which may limit the generalisability of the validation data to some extent. The sample included an unequal although substantial number of women (283) and men (66). However, our use of Bayesian statistics mitigates potential concerns regarding sample size imbalance in gender comparisons by directly estimating the probability of observed effects (Morey, Romeijn, and Rouder 2016; Rouder et al. 2009; Dienes 2011) and accounting for the uncertainty introduced by differing group sizes (Kruschke 2013). Nevertheless, the higher proportion of women may have influenced overall group-level responses, particularly given the observed gender differences. Future research could benefit from a more diverse participant pool to enhance the broader applicability of these findings across different demographics.

In this study, we aimed to create standardised videos with consistent tactile interactions that varied primarily in their hedonic qualities, levels or threat, and perceived arousal. Painful interactions, such as a stab with scissors, were designed to evoke discomfort in the observer (without causing actual harm to the model), while pleasant videos featured gentle, non-threatening touches, such as stroking with a brush. Some neutral scenarios (e.g., a push with a spoon or whisk) may be less naturalistic than other interactions, potentially impacting arousal and threat ratings. We did not collect naturalness ratings, such as those used in the validation of the Socio-Affective Touch Expression Database (SATED; Masson and Beeck 2018), but these could provide additional insights into how contextual relevance shapes touch perception.

In conclusion, the VTD provides a comprehensive resource for investigating how visually presented touch is perceived and processed. By providing a systematically controlled and dynamic database, the VTD facilitates research on both the emotional-affective and sensory dimensions of observed touch, advancing our understanding of visual touch perception.

## CRediT authorship contribution statement

**Sophie Smit:** Conceptualisation, Methodology, Project administration, Data curation, Formal analysis, Visualisation, Writing – original draft, Writing – review & editing. **Anina N. Rich:** Conceptualisation, Methodology, Writing – review & editing, Supervision.

## Additional information

This work was supported by a Commonwealth funded Research Training Program and Macquarie University Research Excellence Scholarship awarded to SS. ANR is supported by an ARC Future Fellowship (FT230100119). Authors declare no competing interests.

## Open practices statement

All data, analysis code, and materials are available at https://osf.io/jvkqa/.

